# Pairing your Sox: Identification of Sox11 partner proteins and interaction domains in the developing neural plate

**DOI:** 10.1101/2020.04.23.057919

**Authors:** Kaela S. Singleton, Pablo Silva-Rodriguez, Elena M. Silva

**Author notes:** Interdisciplinary Program in Neuroscience, Georgetown University Medical Center, Washington D.C., 200057. Regents Hall, 37^th^ and O Street NW, Washington DC 20057. Corresponding author: Elena M. Silva.

## Abstract

Sox11, a member of the SoxC family of transcription factors, has distinct functions at different times in neural development. Studies in mouse, frog, chick and zebrafish show that Sox11 promotes neural fate, neural differentiation, and neuron maturation in the central nervous system. These diverse roles are controlled in part by spatial and temporal-specific protein interactions. However, the partner proteins and Sox11-interaction domains underlying these diverse functions are not well defined. Here, we identify partner proteins and the domains of *Xenopus* Sox11(xSox11) required for protein interaction and function during neurogenesis. Our data show that Sox11 co-localizes and interacts with Pou3f2 and Ngn2 in the anterior neural plate and in early neurons, respectively. We also demonstrate that xSox11 does not interact with Ngn1, a high affinity partner of Sox11 in the mouse cortex, suggesting that Sox11 has species-specific partner proteins. Additionally, we determined that the N-terminus including the HMG domain of xSox11 is necessary for interaction with Pou3f2 and Ngn2, and established a novel role for the N-terminal 46 amino acids in the establishment of placodal progenitors. This is the first identification of partner proteins for *Xenopus* Sox11 and of domains required for partner protein interactions and distinct roles in neurogenesis.

## Introduction

The <u>S</u>ry-related HMG b<u>ox</u> (Sox) family of transcription factors play critical regulatory roles in the development of vertebrates and invertebrates (1, 2). All Sox proteins contain a high mobility group (HMG) domain that binds both partner proteins and DNA in order to alter expression of downstream target genes (3, 4). Sox transcription factors are grouped into eight subfamilies (A-H), based on sequence homology and functional similarity. The SoxB and SoxC subfamily of proteins play essential roles in orchestrating neurogenesis in the central nervous system (CNS). SoxB1 proteins drive neural progenitor specification and maintenance and SoxC proteins promote differentiation and neuron maturation (5–7). Both SoxB and SoxC proteins have multiple, and sometimes opposing, roles during neurogenesis. For example, the SoxB2 protein, Sox21, inhibits neuronal differentiation when over-expressed but is also required at low levels for differentiation (8). Similarly, the SoxB1 protein Sox2, is implicated in embryonic stem cell specification and maintenance, neural progenitor cell competency, and neuron maintenance (9–11). Thus, functional studies in numerous organisms demonstrate that Sox proteins perform unique functions in different neural cell types and at various times during neurogenesis.

Across subfamilies, the HMG domain of Sox proteins is more than 50% identical, resulting in all Sox proteins binding a similar DNA motif with low affinity (12). Interactions with partner proteins are required to facilitate specific and high affinity binding of Sox proteins to DNA regulatory regions. The most commonly identified partners of Sox proteins in neurogenesis are Pou and E-Box proteins (13–15). For example, Sox2 complexes with Pou5f1, Pou3f2 and Ngn2 at various stages in development and drives the expression of different genes in multiple developmental processes (8, 16–18). Sox2 cooperates with Pou5f1 (Oct3/4) to control embryonic stem cell differentiation and later complexes with Pou3f2 (Oct7) to promote neural specification (19–21). Sox proteins can also form homo and heterodimers; Sox2 interacts with Sox21 to promote ectodermal cell fate in stem cells (17). Additionally, Sox2 and Sox21 each bind Ngn2 to promote or inhibit neural differentiation, respectively (8, 22). These data indicate that identifying and characterizing partner protein interactions is essential to understanding the regulation of Sox protein function.

Sox proteins use various domains to complex with partner proteins. While many transcriptional partner proteins interact with the Sox HMG domain (20, 23–27), some interact with multiple domains or a domain outside of the HMG. Sall4 interacts with the C-terminus, N-terminus and HMG domain of Sox2 in embryonic stem cells. Others, like HDAC1, interact with only the C-terminus of Sox2 (28). It has been proposed that protein interactions with domains outside of the HMG serve to stabilize binding of other transcriptional partner proteins (23, 29). The reliance on partner proteins for specificity allow for two levels of regulation of Sox protein function: availability of partner proteins within a given cell or tissue type and the relative affinity of partner protein interactions. Through identification of both Sox-partner proteins and interaction domains, we can reveal how Sox proteins precisely control gene networks during neurogenesis.

In this study, we identify partner proteins and the interaction and functional domains of the SoxC protein, Sox11, during neurogenesis. Work from our lab and others, has shown that Sox11 is involved in neural induction, and neuron differentiation and maturation (30–33). We recently, characterized Sox11 expression and function in the *Xenopus* neural plate and mouse cortex (34). *sox11* expression is found throughout the developing neural plate and cortex, and promotes neuronal maturation in both mouse cortical development and *Xenopus* neurogenesis. Interestingly, we also found that Sox11 is not functionally interchangeable between these two species like many other Sox transcription factors (34, 35). This suggests that Sox11 is essential for neuron formation across species, but the molecular mechanism underlying Sox11 function is not conserved.

To investigate this conundrum and to better understand how *Xenopus* Sox11 drives neurogenesis, we used an open access single-cell sequencing database to characterize expression of Sox11 and potential partner proteins Ngn1, Ngn2, and Pou3f2. We found that *sox11*^*+*^ and *pou3f2*^+^ cells are localized to the anterior neural plate, while *sox11*^+^ and *ngn*^+^ cells are found in early neurons. We also establish that *Xenopus* Sox11 interacts with Ngn2 and Pou3f2 (also known as Brn2), known partners of Sox11 in mouse cortex (31), but does not interact with Ngn1. We went on to identify the domains of Sox11 needed for both partner protein binding and formation of neural progenitors and mature neurons. Collectively, our data show that the first 46 amino acids in the N-terminus of Sox11 is required for robust interaction with Ngn2 and Pou3f2 and formation of neural progenitors in posterior placodes, while the HMG domain of xSox11 is required for both partner protein binding and xSox11 function. Additionally, the C-terminus and transcription activation domain of xSox11 are necessary for formation of neurons in the developing neural plate. These data suggest that Sox11 has species-specific partner proteins to facilitate neuron formation, and that *Xenopus* Sox11 interacts with different partners to promote numerous functions in various regions of the developing neural plate. These are the first partner proteins identified for *Xenopus* Sox11, and the first characterization of Sox11 protein interaction domains and their contribution to neurogenesis.

## RESULTS

### Sox11 and potential partners are co-expressed in distinct cell types of the neural plate

Our previous overexpression studies demonstrated that even though *Xenopus* Sox11 (xSox11) and mouse SOX11 (mSOX11) promote neural differentiation in their respective species, neither promotes differentiation in the other species. One possibility is that xSox11 and mSOX11 utilize different partner proteins and therefore different mechanisms to promote differentiation. Therefore, we asked if xSox11 and mSOX11 interact with the same partner proteins, and focused on three proteins that bind to mSOX11 in the mouse cortex, Ngn1, Ngn2, and Pou3f2 (31).

To determine if the *Xenopus* orthologues of Ngn1, Ngn2, and Pou3f2 are co-expressed with xSox11 in the developing neural plate, we screened an open access database, Jamboree, that contains a time series of single cell transcriptome measurements during *Xenopus* development (36). Our analysis focused on cells in the neural plate marked by *sox2* and *sox3*, the anterior neural plate marked by *nkx2* and *fezf1*, the posterior neural plate marked by *vgll3*, and early neurons marked by *n-tubulin, myt1*, and *prph1* (36). Using this database, we identified cells positive for *sox11* and either *ngn* or *pou3f2*. Due to low sequencing depth, we could not distinguish between *ngn1, ngn2* or *ngn3*. Thus, for the data presented here, we grouped all *ngn*^+^ cells together.

We identified 2,106 *sox11*^+^ cells in the neural plate at stage 12 (late gastrula). At early neurula (stage 13 and 14) there are 924 *sox11*^+^ cells in the anterior neural plate and 1,112 *sox11*^+^ cells in posterior neural plate (Figure 1A, Table S1). In addition, at stage 14, there are 505 *sox11*^+^ cells in early neurons (Figure 1A, Table S1). We examined the number of cells that are *sox11*^+^, *sox11*^+^ and *ngn*^+^, or *sox11*^*+*^ and *pou3f2*^+^, and found that the number of cells coexpressing *sox11, ngn* or *pou3f2* is high at every developmental stage. Of those, only 12 cells in the neural plate and 24 cells in early neurons are positive for all three, *sox11*^+^, *ngn*^+^, and *pou3f2*^+^.

**Figure 1.**
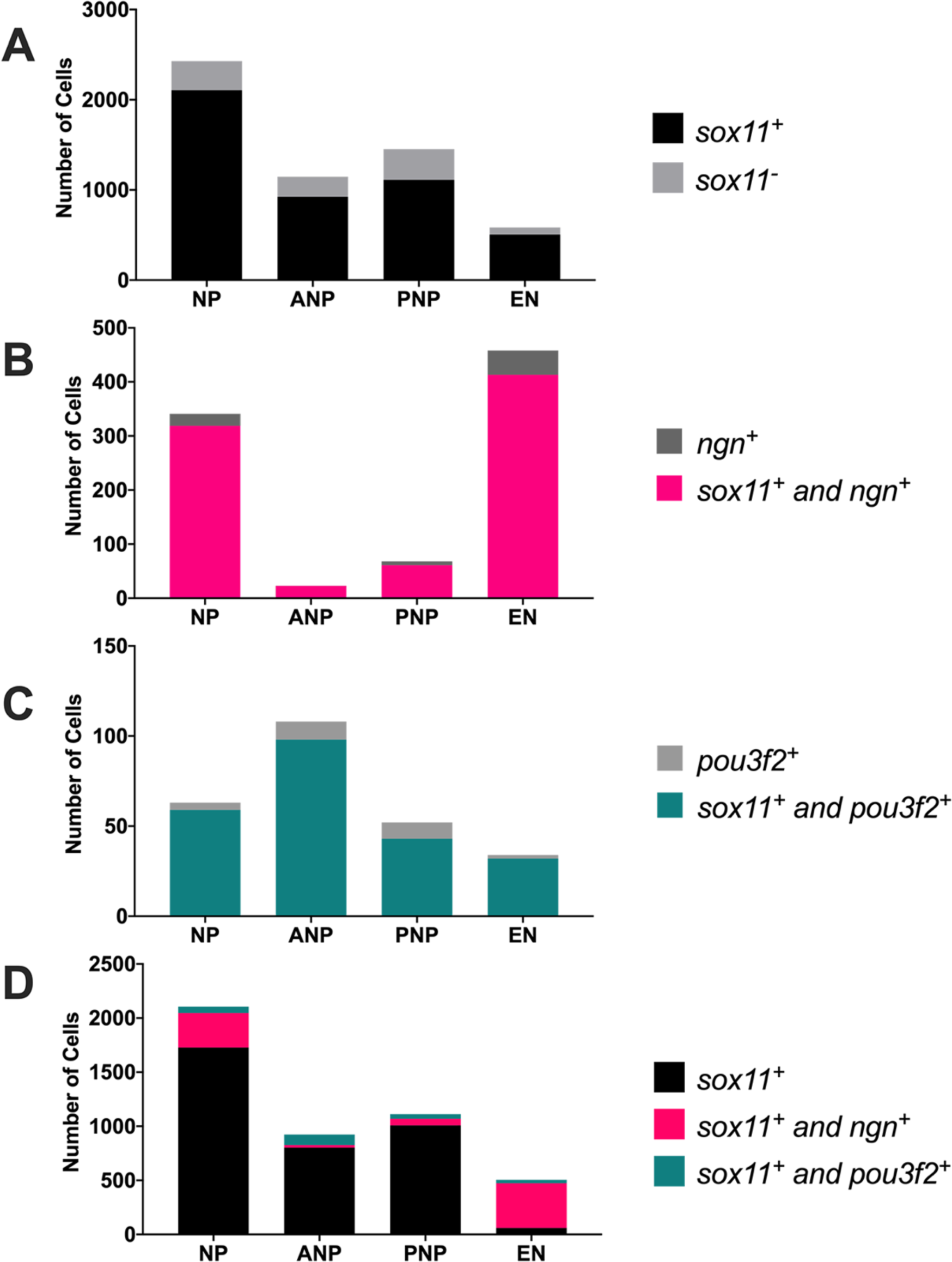
Sox11 and potential partners are co-expressed in distinct cell types of the neural plate. **A** Total number of cells that are *sox11*^*+*^ (black) or *sox11*^*-*^ (gray) within the neural plate (NP) at stage 12, the anterior neural plate (ANP), posterior neural plate (PNP) and early neurons (EN). **B-C** Number of cells across developmental stages that are *ngn+* (dark gray), *ngn+* and *sox11+* (magenta) or *pou3f2+* (light gray) and *pou3f2+* and *sox11+* (teal). **D** Number of *sox11*^*+*^ (black) *sox11*^*+*^ and *ngn*^*+*^ (magenta), *sox11*^*+*^ and *pou3f2*^+^ (teal) cells grouped together.

We show that a majority of *ngn*^+^ cells are *sox11*^+^ (Figure 1B, Table S2); 319 of the 341 *ngn*^+^ cells, in the neural plate are *sox11*^+^, by stage 13/14 all 29 *ngn*^+^ cells in the anterior neural plate are *sox11*^*+*^, and 39 of the 44 *ngn*^*+*^ cells in the posterior neural plate are *sox11*^+^ (Figure 1B). The number of *ngn*^*+*^ cells peaks in early neurons at stage 14; 413 of the 458 *ngn*^+^ cells are *sox11*^+^. Although there are far fewer *pou3f2*^*+*^ cells in late gastrula and early neurula embryos, the majority are also *sox11*^+^ (Figure 1C, Table S3). The highest number of *pou3f2*^*+*^ cells are in the anterior neural plate (108 cells) and 98 are *sox11*^+^. Thus, *sox11* and potential partner proteins are co-expressed in distinct domains of the neural plate (Figure 1D, Table S4). We also confirmed coexpression by performing whole mount *in situ* hybridization (WISH) analysis of stage 12 (late gastrula) and stage 14 (neurula) *Xenopus* embryos demonstrating that *sox11* expression overlaps with *ngn2* an*pou3f2* (Figure S2). In line with our WISH data, previous research shows *pou3f2* expression is not detectable at stage 12 by WISH due to low transcript levels, but is visible at stage 14 in the anterior neural plate (37).

Collectively, these data suggest *sox11* and *ngn* are co-expressed in the correct place and time to promote neuronal differentiation in late gastrula and early neuron formation in neurula *Xenopus* embryos (Figure 1D). Furthermore, *sox11* and *pou3f2* are also co-expressed in the correct place and time to establish the anterior neural plate in early neurula embryos.

### xSox11 partners with Ngn2 and Pou3f2 but not Ngn1

To examine the interactions of xSox11 and potential partner proteins, we generated epitope-tagged expression plasmids: xSox11-FLAG, xPou3f2-HA, xNgn1-HA, and xNgn2-MYC. We performed in vitro translation (IVT) followed by co-immunoprecipitation (co-IP) and western blot analysis (WB) to assay for interactions between xSox11-FLAG and either xPou3f2-HA, xNgn1-HA, or xNgn2-MYC. We show that xSox11-FLAG interacts with both xPou3f2-HA and xNgn2-MYC (Figure 2A-B), but not with xNgn1-HA (Figure 2C). This was surprising because in mouse cortex, mouse NGN1 (mNGN1) is the preferred partner of mouse SOX11 (mSOX11), based on the abundance of the co-immunoprecipitated proteins (31). To establish if xSox11 interacts with mNGN1, we performed co-IP and WB in HEK293 cells. We show that xSox11 does not interact with mNGN1 (Figure 3A), but mSOX11 does (Figure 3B). This suggests that Sox11 has species-specific partner proteins to drive neural differentiation.

**Figure 2.**
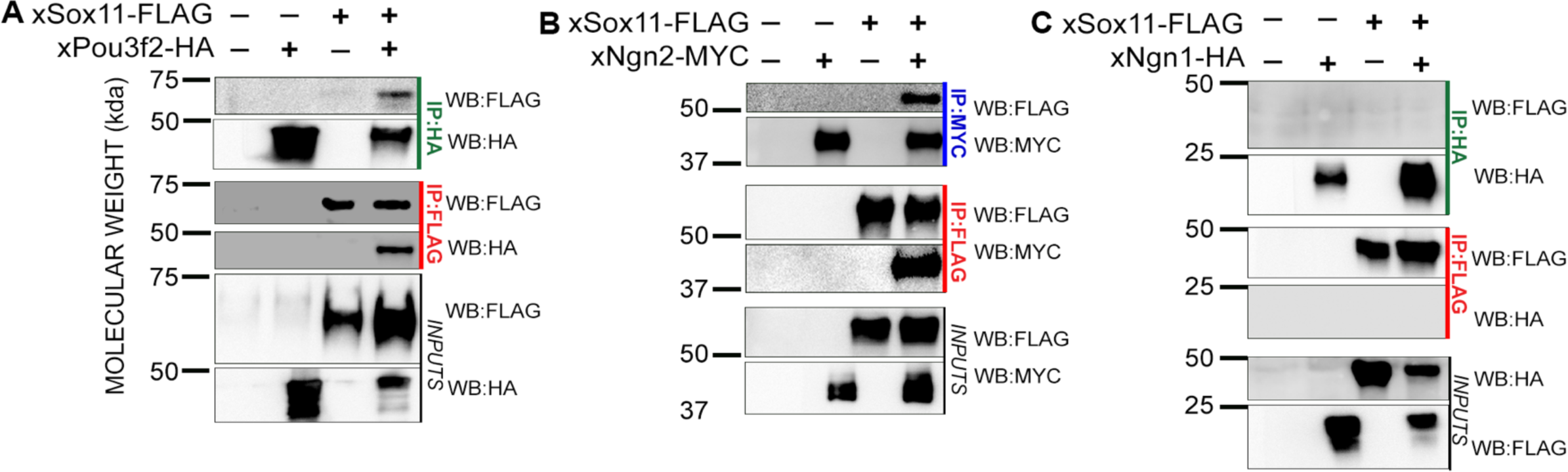
*Xenopus* Sox11 interacts with xPou3f2 and xNgn2, but not xNgn1. **A-D** Immunoprecipitation (IP) of xSox11-FLAG and xPou3f2-HA (A), xNgn2-MYC (B) or xNgn1-HA (C) from *in vitro* translated proteins. Proteins were immunoprecipitated using either FLAG (red), HA (green) or MYC (blue) antibodies. Samples were analyzed by western blot (WB) indicated on the right with FLAG-HRP, MYC-HRP or HA-HRP. Inputs were run as control.

**Figure 3.**
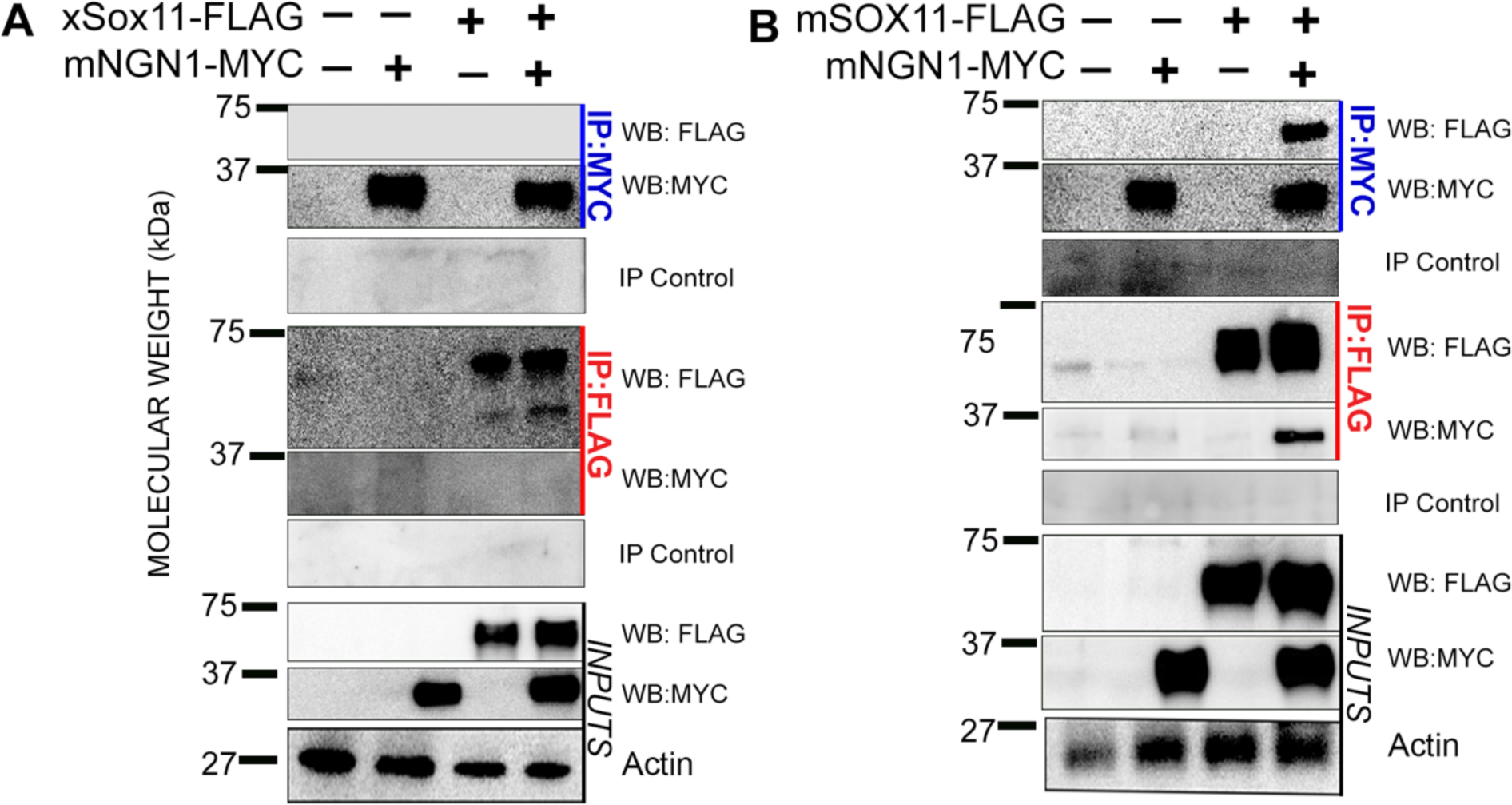
mSOX11 interacts with mNGN1, but xSox11 does not. **A-B** Immunoprecipitation (IP) of xSox11-FLAG (A), mouse SOX11-FLAG (mSOX11-FLAG, B), with mouse NGN1-MYC (mNGN1-MYC) using either FLAG (red) or MYC (blue) antibodies in HEK293 cells. Samples were analyzed using western blot (WB) with either anti-FLAG-HRP or anti-MYC-HRP. Inputs, actin, and IgG (IP Control) were run as control.

### Sox11 N-terminus is necessary for protein-protein interactions

To investigate which domains of Sox11 are necessary for partner protein binding, we generated constructs that express modified xSox11-FLAG protein: ΔN46-xSox11-FLAG that removes 46 amino acids upstream of the HMG domain, ΔHMG-xSox11-FLAG that removes the entire 72 amino acid HMG domain, and the ΔCterm-xSox11-FLAG that removes 265 amino acids and contains only the N-terminus and HMG of Sox11 (Figure 4A).

**Figure 4.**
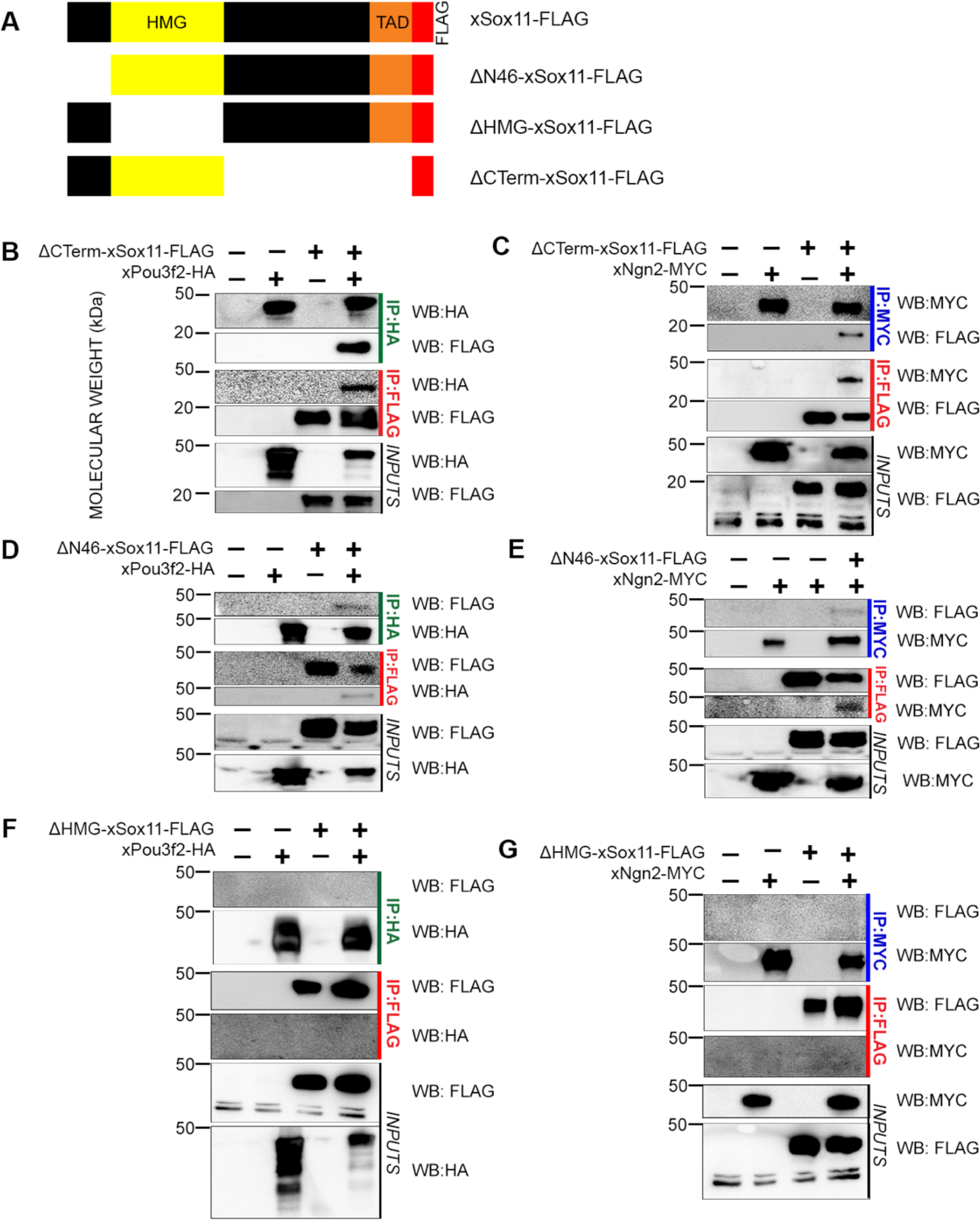
Sox11 N-terminus is essential for protein-protein interactions. **A** Schematic of xSox11domains including HMG domain (yellow), transactivation domain (orange) and FLAG tag (red) along with xSox11 deletion constructs. ΔN46-xSox11-FLAG that removes 46 amino acids upstream of the HMG domain, ΔHMG-xSox11-FLAG that removes the 72 amino acid HMG domain, and the ΔCterm-xSox11-FLAG that removes 265 amino acids and contains only the N-terminus and HMG domain. **C-G** Immunoprecipitation (IP) of ΔCterm-xSox11-FLAG, ΔN46-xSox11-FLAG or ΔHMG-xSox11-FLAG with xPou3f2-HA or xNgn2-MYC. Proteins were immunoprecipitated using either FLAG (red), HA (green), or MYC (blue) antibodies. Samples were analyzed by WB with anti-FLAG-HRP, anti-HA-HRP, or anti-MYC-HRP. Inputs were ran as control.

To map the protein interaction domains, we performed co-IP of the modified xSox11 proteins and xNgn2 and xPou3f2 using FLAG, HA or MYC antibodies and analyzed samples by WB. Interestingly, the loss of the xSox11 C-terminus does not affect binding of either xPou3f2-HA or xNgn2-MYC (Figure 4B-C), and suggests that the C-terminus is not necessary for Sox11interactions with these two proteins. We next examined how the loss of the N46 xSox11 domain altered interactions with xPou3f2-HA and xNgn2-MYC. Our results show that removal of the N46 domain significantly weakens the interaction with both partners (Figure 4D-E). Lastly, removal of the HMG domain prohibits interaction with either partner protein (Figure 4F-G). These data reveal that the short 46 amino acid N-terminus of Sox11 is required for strong partner protein interactions and the HMG domain is necessary for partner protein binding.

### Sox11 C-terminus is required for mature neuron formation

Our previous work demonstrated that overexpression of xSox11 increases mature neurons marked by *n-tubulin* (*n-tub*), and increases neural progenitors marked by *sox3* (34). Furthermore, we showed that loss of xSox11 reduces mature neurons and increases *sox3* (34). To determine the functional significance of xSox11 domains, we analyzed the effect of our mutant xSox11 constructs on neurogenesis (Figure 4A). To do this, we injected mRNA from each construct in one cell of a two-cell blastomere embryo, allowed embryos to develop to neurula stage, and assayed by WISH for changes in *n-tub* and *sox3*. For domains that are necessary for xSox11 function, we expect to see no change or a decrease in *n-tub* and/or *sox3* similar to our xSox11 knockout data, and for domains that are not necessary for xSox11 function, we expect to see an increase in *n-tub* and/or *sox3* in line with our previously published data (34).

Our data reveal that with overexpression of *Δcterm*-*xsox11, sox3* expression remained the same, but *n-tub* expression was reduced. This result indicates that the ΔCterm-xSox11 functions as a dominant negative and like the MO (34) and reduces mature neuron formation. Overexpression of *Δn46*-*xsox11* leads to ectopic expression of *n-tub* in the neural plate and loss of *sox3* expression in placodes (Figure 5). These data show that the 46 N-terminal amino acids are not required for promotion of neural differentiation but is required for placode development. Here the ΔN46-sox11 interferes with placode formation and thus functions as a dominant negative.

**Figure 5.**
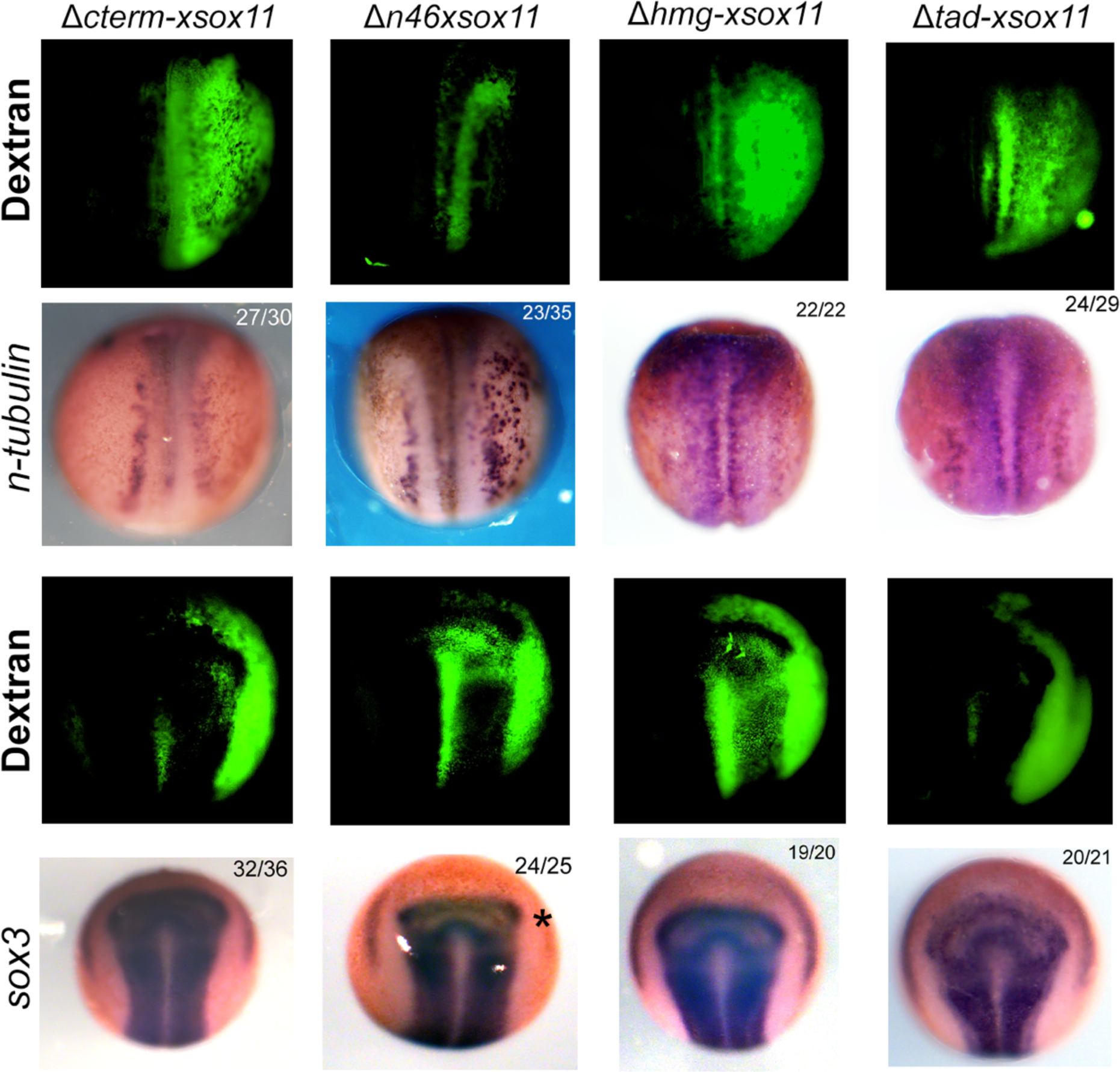
Sox11 C-terminus controls mature neuron formation. WISH of neurula (stage 15) embryos injected in one of two cells (dorsal view, anterior to the top) with Dextran (green) and either *Δcterm-xsox11, Δn46-xsox11, Δhmg-xsox11*, or *Δtad-xsox11* mRNA. Embryos were then analyzed for expression of *n-tubulin* for mature neurons or *sox3* for neural progenitors. Asterisk shows reduction in placodal progenitors. Numbers in the upper right of each image denote the number of embryos with the phenotype over the total of embryos analyzed.

Next, we overexpressed *Δhmg-xsox11*. Our data show that there is no difference for either *n-tub* or *sox3* expression, which we predicted given that Sox transcription factors cannot bind DNA or complex with partners when the HMG is removed (38). We also investigated the role of the 48 amino acid trans-activation domain (TAD) located in the C-terminus of Sox11 (39, 40). We generated a *Δtad-xsox11* mutant and assayed for its ability to produce protein (Figure S4) and performed the same overexpression experiments analyzing *n-tub* and *sox3* (Figure 5). We found that without the TAD, xSox11 shows a slight decrease in *n-tub* and no change in *sox3* expression. This result is in line with previous studies that show the TAD is essential for Sox11 function *in vitro* (41). Importantly, this is the first time domains of Sox11 have been characterized *in vivo* to determine their contribution to neurogenesis. Together, our data show that HMG domain is necessary for xSox11 function during neuron formation. Additionally, our data reveal that the N46 domain of xSox11 is necessary for development of neural progenitors in the placodes, and that the C-terminus of xSox11, including the TAD, is necessary for mature neuron formation.

## Discussion

Sox transcription factors cooperate with region-specific partner proteins to regulate down-stream targets and orchestrate neurogenesis. To identify and characterize Sox11-partner protein interactions essential to neurogenesis, we tested whether Sox11 partner proteins are conserved between mouse and *Xenopus* and identified the domains of Sox11 necessary for protein interaction and function. Previous studies identified Ngn1, Ngn2, and Pou3f2 as binding partners of Sox11 in mouse cortex and in cos9 cells (31, 40). We aimed to establish if these interactions also occur in the developing *Xenopus* neural plate. Our co-expression analysis, using a single cell sequencing database (36), revealed that xSox11 is expressed in the same cells as Ngn1, Ngn2 and Pou3f2 at different times in early neural development. *sox11* and *ngn* are co-expressed predominantly in early neurons, and *sox11* and *pou3f2* cells are abundant in the anterior neural plate (Figure 1). Thus, xSox11 and these candidate partner proteins are expressed at the same time and place to interact. Through a series of co-IP experiments, we show that xSox11 interacts with Ngn2 and Pou3f2, but not Ngn1 (Figure 2). Importantly, the role of Ngn2 in neurogenesis has been well characterized and complements Sox11 function; Ngn2 governs formation of the first neurons generated in the CNS (42), and Sox11 drives the formation of early born neurons (43). Thus, our data support the relationship between xSox11 and Ngn2 as essential regulators of early neuron formation.

We also show that xSox11 interacts with xPou3f2 (also known as Brn2 or Oct7), however, this relationship is not well defined in *Xenopus* neural plate development and has various expression patterns and functions across species. Pou3f2 is robustly expressed in stage 14 neurulae but is not detectable in stage 12 gastrulae (Figure SA1) (37). In rat neural precursor cells, Pou3f2 co-localizes with progenitor cell markers and is downregulated upon differentiation (44). However, in mouse cortex, Pou3f2 expression is restricted to upper layer neurons, and loss of both Pou3f2 and family member Pou3f1 results in a decrease in neuronal migration, layer production and neurogenesis (45–47). Thus, Pou3f2 plays critical, but disparate, roles in neurogenesis. Despite this, few studies have investigated the relationship between Sox11 and Pou3f2. Previous reporter gene studies show that Pou3f2 inhibits mouse Sox11 transactivation (31). Since neural differentiation is delayed in the anterior neural plate where *pou3f2* and *xsox11* are co-expressed, our data supports the idea that Pou3f2 antagonizes the ability of Sox11 to promote neurogenesis (Figure 2) (48). Collectively, these data suggest Pou3f2 inhibits Sox11 function to delay neuronal differentiation.

Our studies reveal that xSox11 does not interact with either *Xenopus* or mouse Ngn1, suggesting that the partner proteins of Sox11 vary across species (Figure 3). This was a surprise since mouse Ngn1 is the preferred partner for Sox11 in mouse cortical cells, while Sox4, another SoxC family member, preferentially complexes with Ngn2 (31). One interesting possibility, is that Sox11 and Sox4 evolved to interact with different Ngn proteins to allow for the expansion of the mammalian cortex. Sox4 has been partially characterized in *Xenopus* but only in the eye (49). Thus, future studies on the role of Sox4 in the neural plate and tube of *Xenopus* could address these questions. A second possibility is that post translational modification (PTM) or a mediator protein in neural tissue is required for Ngn1 to interact with xSox11. There is precedent for PTMs of mouse Sox11 in developing mouse retina and hippocampus (50, 51). To our knowledge, no studies have investigated PTMs of xSox11 or Ngn1 during neurogenesis. Importantly, our studies were conducted *in vitro* and did not allow for PTMs of either protein. Mediator proteins have been shown to complex with Ngn1 and other Sox proteins to stabilize reactions and alter function (23). Specifically, when Ngn1 complexes with CBP/p300, it facilitates an interaction with Smad1 to inhibit glial cell differentiation and promote neurogenesis (52). Additionally, Sox2 and Sox6 have been shown to use co-activators and repressors (53–55), and Sox4 has also been shown to require both cytokine interaction and PDZ class proteins to regulate receptor expression (56). These possibilities can be explored in future studies.

In addition to identifying partner proteins of xSox11, we identified the interaction domains. We show that the HMG domain of Sox11 is necessary for partner protein binding, and the first 46 amino acids of the N-terminus (N46) are essential for strong partner interactions (Figure 4). The HMG domain has been shown to be essential for protein interaction in other Sox proteins. For example, SoxE proteins (Sox8, Sox9, Sox10) require the C-terminal tail of the HMG to complex with partners (57, 58). However, in other cases, domains outside of the HMG are required for partner protein interactions. The B-homology domain and C-terminus of Sox2 is required for partner interactions in stem cells (28) and Sox18 binds to MEF2C in endothelial cells through its C-terminal domain (59). Thus, similar to other Sox proteins, Sox11 uses a domain outside of the HMG, to complex with partners, and this likely contributes to its binding specificity (15).

Our previous work demonstrated that overexpression of xSox11 increases both mature neurons, and neural progenitors (34). Additionally, we showed that knockdown using a morpholino (MO) of xSox11 results in a decrease in mature neurons, and leads to an increase in neural progenitors, potentially due to progenitors proliferating rather than differentiating. Together, these data establish xSox11 as a critical protein during early neurogenesis. Our analysis of the functional significance of Sox11 domains revealed embryo phenotypes that support roles for protein domains outside of the HMG in protein interaction. Overexpression of ΔCterm-xSox11 mimics the knockdown phenotype: decreased neuron formation as marked by *n-tub* expression (34). This is likely due to ΔCterm-xSox11 functioning as a dominant negative by binding partner proteins in the developing neural plate (Figure 4B-C) and preventing endogenous Sox11 from interacting with these proteins. Since ΔCterm-xSox11 lacks the TAD, it is unable to activate target genes. Interestingly, overexpression of ΔCterm-xSox11 does not affect neural progenitors in the same way as xSox11 knockdown. The reasons for this are unclear, but since the increase in progenitors is slight in the Sox11 knockdown embryo, this could simply be due to ΔCterm-xSox11 being less effective than the MO. Additionally, we show that excess ΔN46-xSox11 functions similarly to overexpression of xSox11 in embryos and causes an increase in mature neurons. Our binding studies showed that ΔN46-xSox11 interacts with partner proteins (Ngn2), and since it has an intact C-terminus and TAD, can still activate Sox11 downstream targets. We also show that overexpression of ΔN46-xSox11 decreases neural progenitors in the placodes (Figure 5). To further investigate this result, we used the single cell sequencing database (36). We identified posterior placodal cells as those expressing *pax8, six1*, and *sox9* and found 487 *sox11*^*+*^ cells out of 618 total cells, supporting the role for Sox11 in placodal development as previously described (60, 61). We also confirmed that neither *pou3f2* nor *ngn* is expressed in posterior placode. Thus, Sox11 likely works with an unknown placodal partner protein to regulate posterior placodal development. Together, these data show that the C-terminus of Sox11 is essential for neuron formation, and that further work must be done to elucidate Sox11 partner proteins in placodal progenitors.

Our overexpression studies demonstrate that ΔHMG-xSox11 has no detectable effect on mature neurons or neural progenitors, while overexpression of ΔTAD-xSox11 shows a slight decrease in mature neurons and no change in neural progenitors (Figure 5). These results are in line with previous studies that demonstrate the importance of the HMG and TAD domains in other Sox proteins. Numerous studies have shown that the HMG domain is essential to both DNA and partner protein binding (38, 62). Additionally, our data underscore the significance of the serine-rich TAD of Sox proteins. Importantly, the TAD of Sox11 was first shown to be essential in oligodendrocytes and characterized further in cos9 cells (39, 63). Sox11 is known as the most potent transactivator of the SoxC family, and has even been shown to be more potent than Sox2 (40).

In conclusions, here we uncover several critical features of Sox11, a protein necessary for neurogenesis. First, xSox11 is co-expressed with xPou3f2 and xNgn2 in the anterior neural plate and early neurons, respectively. Second, Sox11 partner proteins are not conserved across species leading to the enticing possibility that changes in SoxC proteins evolved to enable expansion of the cortex in mammals. Third, the HMG domain and the first 46 amino acids in the N-terminus are necessary for robust partner protein interactions, whereas the C-terminus plays no role in the binding of Pou3f2 and Ngn2. Lastly, we show that the C-terminus of Sox11, and specifically, the TAD is required for promoting neuron formation and progenitors, and the N46 domain of xSox11 is essential in posterior placodal development. To our knowledge, these are the first partner proteins identified for *Xenopus* Sox11 and the first identification of Sox11 domains essential for protein interaction and neuron formation in the developing neural plate.

## Experimental Procedures

### Plasmids

pCS2+ (used as control in HEK experiments), mNgn2-MYC (gift from Qiang Lu), xPou3f2-HA, xNgn1-HA (HA versions generated by GeneWiz), xNgn2-MYC (gift from Sally Moody), mSox11-FLAG (gift from Maria Donoghue) and xSox11-FLAG were used. Sox11-FLAG mutants were created by site-directed mutagenesis via polymerase chain reaction with 2XPfu DNA polymerase (Agilent). Primers can be found in Table 1. Each plasmid expressed the correct sized protein as determined by immunoprecipitation (IP) and western blot (WB) analysis (Figure S2-S4).

**Table 1.**
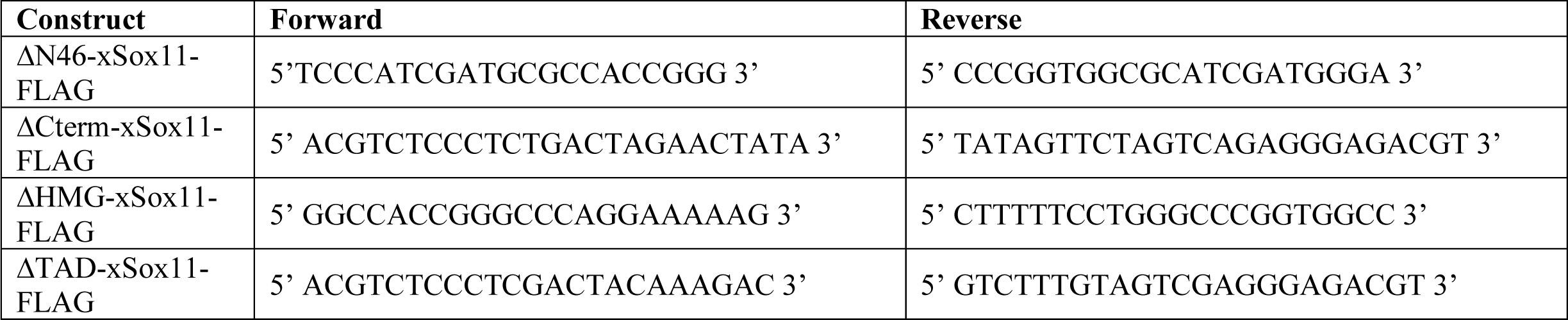
Primers for xSox11 deletion constructs.

### Identifying Cells using *Xenopus* time Series Database

Single cell transcriptome measurements were taken from Xenopus Jamboree database (klein-tools.hms.harvard.edu). Cells in the following categories were analyzed: Neural Plate (Stage 12), Anterior Neural Plate (Stage 13 and 14), Chordal (Posterior) Neural Plate (Stage 13 and 14), and Early Neuron (Stage 13 and 14).

### *Xenopus* animal usage and embryo manipulation

All frog use and care were in accordance with federal and institutional guidelines, particularly Georgetown University’s Institutional Animal Care and Use Committee protocols 13-016-100085. *Xenopus laevis* embryos were obtained using standard methods (64, 65) and staged according to the Nieuwkoop & Faber staging system.

### Whole Mount In situ hybridization

Whole mount in situ hybridization (WISH) was performed as previously described (66, 67) with these changes: length of pre-hybridization step was increased to overnight and the RNAse treatment step was eliminated. Digoxygenin labeled probes were generated from plasmids produced by the Silva laboratory.

### Frog microinjection

mRNA used for injections were made in vitro using the mMESSAGE mMACHINE® Transcription Kit (Life Technologies). Embryos were injected with mRNA (1.2ng) and 1:5 Dextran (tracer, ThermoFisher) in one cell of two-cell stage embryos to over-express *sox11* or *sox11* deletion constructs. Embryos were cultured until neurula stage (st. 14) in 1/3 MMR at 17 °C and fixed in Bouins fixative. Embryos were visualized under 488 wavelength fluorescence to identify the injected side following whole mount in situ hybridization experiment.

### In vitro translation (IVT) assay

TNT® SP6 High-Yield Wheat Germ Protein Expression System (Promega) was used to perform in vitro translation assay. Protein products were denatured at 100°C for 10 minutes and separated on SDS-PAGE gel. Antibodies for anti-FLAG-HRP (Sigma, 1:2500), anti-HA-HRP (Roche, 1:2500), anti-cMyc-HRP (Abcam, 1:2500), were used to confirm expression.

### HEK cells maintenance and transfection

HEK293 cells were grown in DMEM (Dulbecco’s Modified Eagle Medium) supplemented with 10% fetal bovine serum and 1% penicillin/streptomycin, and maintained at 37 degrees Celsius with 5% CO2 in a humidified environment. Cells were passaged at ∼70% confluency by brief trypsinization at room temperature, followed by serum inactivation and plating in fresh growth media. HEK cells were transfected at 50-60% confluent in complete growth medium, using X-tremeGENE (Roche), according to manufacturer’s instructions. For one well of a 6-well dish, 2µg total DNA was transfected at 1:1 DNA:reagent ratio, where 0.5 µg of control plus 1.5µg/well of either *Xenopus* or mouse Sox11-FLAG, and/or Ngn1 expression plasmids was added to total 2 µg. Transfection cocktail was added directly to the well and left until cell lysis.

### Co-immunoprecipitation for IVT Protein

*In vitro* translation (IVT) with 500 ng of mRNA was performed using the TnT SP6 High-Yield Protein Expression System (Promega) to synthesize tagged proteins. 66% (20µl) of this reaction was subjected to co-IP was while 33% (10µl) was saved as input. co-IP for IVT was performed as previously described with the following changes (8). For each co-IP, 10 µl reaction mix was mixed with 290µl IP buffer (1% NP-40, 50 mM Tris-HCl, pH 8.0, 150 mM NaCl) containing protease inhibitor cocktail tablet (Roche) and 2 µg/ml of anti-FLAG (Sigma), anti-HA (Cell Signaling), or anti-MYC (Cell Signaling) and incubated for one hour on ice. 25µl Dyna Protein G magnetic beads (ThermoFisher) were added and the reaction was incubated at 4°C overnight on an orbital mixer. The next day, beads were washed three times in cold IP buffer. Buffer was removed and resuspended in 2X SDS sample buffer with DTT and boiled at 100°C for 10 minutes. 12 µl of IP sample and 2.5 µl of input (also boiled in 2X Laemmli Sample buffer with DTT) were separated via SDS-PAGE on 12% pre-cast gels (Bio-rad).

### Co-immunoprecipitation for HEK293 Lysate

Transfected HEK cells were lysed in cold immunoprecipitation buffer (1% NP-40, 50 mM Tris-HCl, pH 8.0, 150 mM NaCl) with protease inhibitors (Roche), and centrifuged for 10 minutes and 3,000 rpm at 4°C. Supernatant was collected and measured via Bradford Protein Assay (Bio-rad). 200µg total protein were used in co-IP assays and 5µl of lysate was used as input. For co-immunoprecipitation, Dyna Protein G magnetic beads (ThermoFisher) were washed in IP buffer and incubated with primary antibody (rabbit anti-HA, anti-myc, CST or mouse anti-flag, Sigma) or normal IgG as control (mouse IgG, rabbit IgG, ThermoFisher) at 1:100 dilution for ≥1hr, then rinsed in IP buffer. 200µg total protein was incubated with prepared magnetics beads in IP buffer overnight at 4°C. Samples were washed in cold IP buffer before being eluted in 2X Laemmli sample buffer with DTT, boiled, and separated via SDS-PAGE on 12% pre-cast gels (Bio-rad).

### Western Blots

Western blots were performed using the Bio-Rad Mini TransBlot and TransBlot-Turbo Transfer System, with the Bio-Rad PVDF Transfer Kit. Briefly, samples were loaded onto 12% Tris-glycine SDS polyacrylamide gels (Bio-Rad), resolved by electrophoresis, transferred to PVDF, and blocked in 5% non-fat dry milk in TBST (TBS+ 0.1% Tween-20) for one hour at room temperature. Following block, blots were incubated with anti-FLAG-HRP (Sigma, 1:2500), anti-HA-HRP (Roche, 1:2500), anti-myc-HRP (Santa Cruz, 1:2500), anti-beta actin (Sigma, 1:5000) primary antibodies overnight at 4°C, washed three times with TBST, then incubated with Pierce ECL Plus chemiluminescent substrate (ThermoFisher) and imaged using ImaqeQuant LAS-4000 mini digital imager (GE Healthcare) and visualized for no more than 5 minutes. Input blots were run as control for each experiment, and actin was run as a loading control for HEK experiments.

## Data Availability

All data are contained within the article.

## Acknowledgments

We thank members of the Silva lab for scientific editing of the manuscript.

## Funding and additional information

This work was supported in part by NIH grant NS078741. KSS is supported by the Center of Neural Injury and Plasticity (5T32NS041218) and NINDS Pre-doctoral to postdoctoral advancement in neuroscience grant (F99NS108539). PSR received support from the Fulbright Foundation.

